# An Insoluble *De Novo* Protein Enables Survival of *E. coli* by Co-precipitating with a Gene Repressor

**DOI:** 10.64898/2026.01.02.697383

**Authors:** Guanyu Liao, Sha Tao, Jessica L. Dessau, Yejin Bann, Michael H. Hecht

## Abstract

*De novo* proteins that share no ancestry with natural sequences can serve as additions to the evolved proteomes of living cells. Upon expression in cells, these novel proteins can provide biological functions that alter cell viability and growth. To isolate such proteins, we searched a combinatorial library of novel sequences by selecting for sequences that sustain the growth of *E. coli* under conditions where the recipient cell would otherwise be inviable. This led to the discovery of *Resc4 (Rescuer 4),* a *de novo* protein that sustains growth on minimal medium of an *E. coli* strain harboring a lethal deletion of *metC,* which encodes cystathionine β-lyase, an essential enzyme in the biosynthesis of methionine. Surprisingly, despite its ability to rescue the deletion of a biosynthetic enzyme, *Resc4* is insoluble. Nonetheless, *Resc4* sustains the growth of 11*metC* cells by upregulating expression of *metB*, which encodes a different enzyme, cystathionine γ-synthase, which has a moonlighting activity that compensates for the deleted activity encoded by *metC.* Proteomic analysis revealed that *Resc4* co-precipitates MetJ, the repressor of the methionine biosynthesis operon. Precipitation of MetJ leads to overproduction of cystathionine γ-synthase, thereby allowing it to rescue the deletion of *metC*. These results, taken together with previous findings on other *de novo* proteins, demonstrate that novel proteins added to a cell’s proteome can perform life sustaining functions, and may shed light on *de novo* gene birth – both in synthetic biology and in natural evolution.

## Introduction

An important goal of synthetic biology is to reconstruct life using genes and proteins unrelated to those that evolved in nature. Toward this goal, *de novo* sequence libraries have been screened for life-sustaining sequences. We previously described several *de novo* proteins that provide life-sustaining functions in *E. coli* under otherwise lethal conditions (1–3). These novel sequences can rescue *E. coli* using a range of different mechanisms. For example, Syn-F4 is a binary pattered 4-helix bundle that rescues the Δ*fes* strain of *E. coli* in iron-deficient media (4). In this case the rescue is direct: Syn-F4 is a catalytically active enzyme that replaces the function of the deleted enzyme, ferric-enterobactin esterase, encoded by *fes* (5, 6).

In other cases, rescue can be indirect, and a *de novo* protein can sustain cell viability by altering gene regulation. For example, we previously reported that the *de novo* protein SynSerB3 rescues the deletion of phosphoserine phosphatase in Δ*serB* cells by upregulating the endogenous enzyme histodinol-phosphatase (encoded by HisB), which has a low level of promiscuous phosphatase activity (7). In another example, deletion of citrate synthase (Δ*gltA*) was rescued by SynGltA by upregulation of the promiscuous enzyme encoded by PrpC, which has a low level of citrate synthase activity (8).

The Laub lab has also shown that biologically significant activities can be isolated from non-biological sequences: By screening libraries of random sequences, Frumkin *et al.* discovered novel proteins that rescue the growth of *E. coli* in otherwise lethal conditions by either modulating protein homeostasis (9) or facilitating resistance to phage (10).

Building on previous successes using our *de novo* 4-helix bundles to provide life-sustaining functions, we set out to screen libraries that might rescue a range of different gene deletions. Specifically, with the goal of discovering additional *de novo* enzymes, we constructed libraries of sequences in which the catalytic site of our *de novo* enterobactin esterase, Syn-F4, was semi-randomized. Unexpectedly, however, screens and selections of this library in several auxotrophic strains of *E. coli* led to the isolation of novel sequences containing frameshifts and stop codons that altered the sequences and properties of the original Syn-F4 protein. Interestingly, several of these novel sequences encode short proteins that are insoluble but nonetheless provide biological functions capable of sustaining the growth of otherwise moribund cells.

Here we describe the *de novo* protein *Resc4*, which was isolated by screening a library of novel sequences for the ability to rescue Δ*metC* cells on minimal medium, wherein methionine biosynthesis is essential for survival. *Resc4* is unusual in several respects: (i) It contains an unexpected frameshift. (ii) It contains an early stop codon, leading to a short (∼84 residue) polypeptide. (iii) *Resc4* is insoluble and enzymatically inactive. (iv) Nonetheless, *Resc4* rescues Δ*metC* cells on minimal medium by enhancing expression of another endogenous enzyme – cystathionine γ-synthase – which has a promiscuous activity capable of rescuing the deletion of *metC*. (v) Genetic, biochemical, and proteomic analysis revealed that *Resc4* accomplishes this unexpected rescue by co-precipitating the MetJ protein, which is a repressor of the methionine biosynthesis operon. With the removal of MetJ from the soluble milieu, the methionine biosynthesis operon is turned up, thereby leading to over expression of the promiscuous cystathionine γ-synthase (MetB) enzyme.

These results demonstrate that novel proteins with no biological ancestry can provide life sustaining functions using unprecedented and unexpected mechanisms.

## Results

### Constructing a protein library based on a de novo enzyme

Towards the goal of discovering novel proteins with life-sustaining catalytic activities, we constructed a library of sequences based on the first *de novo* life-sustaining enzyme, Syn-F4, which rescues Δ*fes* in iron-deficient media by hydrolyzing ferric-enterobactin (5). Specifically, we used degenerate codons to semi-randomize the catalytic residues in Syn-F4 and residues surrounding the active site (**Supplementary Figure 1**).

The crystal structure of Syn-F4 demonstrated that the sequence assembles into a domain-matched homodimer, where the N- and C-termini of both subunits reside on the same end of the helical bundle (6). Inspired by Dayhoff’s hypothesis, which suggests that highly active enzymes can emerge through gene duplication followed by asymmetric diversification (11, 12), we inserted a flexible linker to join the two monomer subunits in the Syn-F4 library, thereby creating a library of asymmetric heterodimers containing 10^6^-10^7^ different sequences (**Supplementary Figure 1**). Since this library was based on the sequence of Syn-F4, the sequences in this library share no ancestry or sequence homology with natural proteins.

### Screening the library for life-sustaining sequences

We searched the library of *de novo* sequences for proteins that rescue the growth of *E. coli* strains with gene deletions that prevent growth on minimal media. This was accomplished by transforming auxotrophic *E. coli* cells with plasmids expressing genes that encode sequences from the library of putative heterodimers. Transformed cells were incubated on selective minimal media for 10 days or until colonies were observed (**Supplementary Figure 2**).

After testing a range of auxotrophic strains with gene deletions in various biosynthetic pathways (**Supplementary Table 1**), we found three strains – Δ*fes*, Δ*ilvA* and Δ*metC* – that were rescued by sequences from the library. While rescuers of Δ*fes* and Δ*ilvA* were described in our earlier work (2, 4, 5), this was the first time we isolated novel sequences that rescue Δ*metC* (**Figure 1A, B**). *metC* encodes cystathionine β-lyase, a key enzyme in the pathway of methionine biosynthesis (13, 14). Several different sequences produced colonies on minimal media within five days. Unexpectedly, however, all of the rescue sequences contained frame-shift insertions that alter every amino acid starting between residue 20 and 40 (**Figure 1A**). Because of these frame shifts, all the rescue sequences have a premature stop codon after residue 84 or 85. In addition, all the frame-shifted rescue proteins are rich in proline and arginine (**Figure 1A**), which likely prevents them from folding into native-like globular structures.

**Figure 1.**
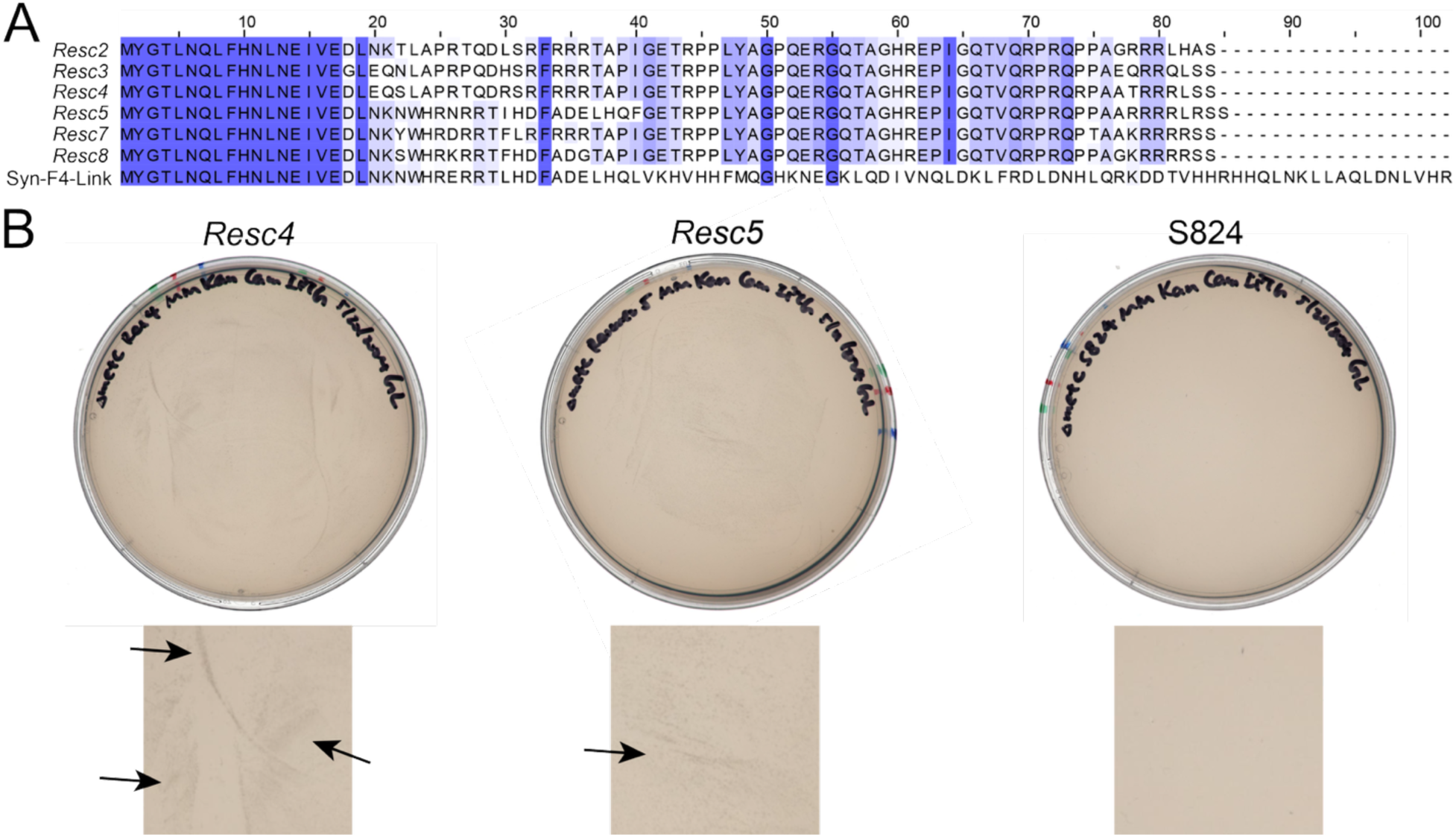
Δ*metC* rescuers enable Δ*metC* to grow on minimal media. (A) Amino acid sequences of frame-shifted Δ*metC* rescuers. The background of the text color represents degree of conservation in the multiple sequence alignment. Dark blue background represents high conservation, light blue represents medium conservation, and white background represents no conservation. Only the first 102 amino acids of Syn-F4-Link sequence are shown. Frameshifts occur between residue 20 to residue 40. (B) Selection plates (top) and zoomed-in views (bottom) for rescue of Δ*metC* cells by *Resc4*, *Resc5* and S842 (negative control). Smears in the zoomed in views (arrows) indicate growth.

Because of the unexpected frame-shifts and truncations, we considered the possibility that novel mRNAs, rather than translated protein sequences, might be responsible for the observed rescue. To test this possibility, we deleted the methionine start codon in the rescuer sequences. The sequences in which the start codon was deleted failed to rescue Δ*metC*, indicating that the protein, not the mRNA, was the functional biomolecule for Δ*metC* rescue (**Supplementary Figure 3**).

### Resc4 cannot catalyze the reaction performed by the natural MetC enzyme

To elucidate the mechanism by which *de novo* sequences rescue Δ*metC*, we first tested whether these proteins could catalyze the same reaction as the natural enzyme, cystathionine β-lyase, encoded by MetC (**Figure 2A**). To probe for activity, we picked a rescuer protein, *Rescuer 4* (*Resc4*), and attempted to purify it. However, *Resc4* was insoluble. Therefore, we added a His_6_-SUMO tag at the N-terminal of the sequence to enhance solubility and facilitate protein purification. We purified the SUMO-tagged *Resc4* in a denaturing environment (8 M urea) and solubilized it by stepwise dialysis into a native buffer (**Figure 2B**).

**Figure 2.**
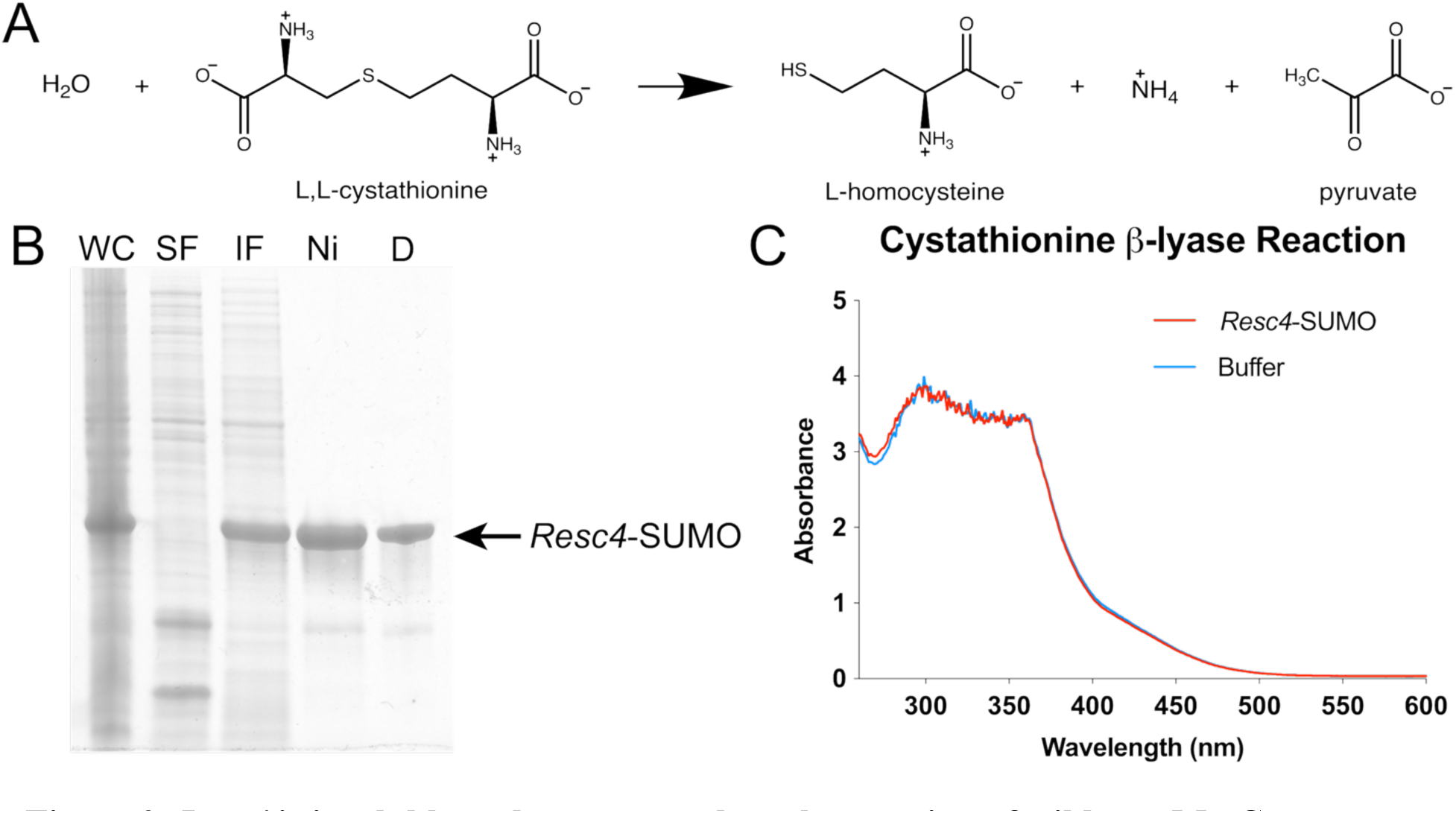
*Resc4* is insoluble and cannot catalyze the reaction of wildtype MetC. (A) Cystathionine β-lyase reaction catalyzed by wildtype MetC. (B) SDS-PAGE gel for purification of *Resc4* with N-terminal His_6_-SUMO tag (*Resc4*-SUMO). WC: whole cell. SF: soluble fraction. IF: insoluble fraction. Ni: Ni-NTA elution fraction. D: soluble protein after dialysis of Ni-NTA elution fraction. (C) UV-vis absorbance of L,L-cystathionine β-lyase reaction. Purified and solubilized *Resc4*-SUMO or buffer control (no protein added) was incubated with PLP and cystathionine at 37 °C for 24 hours. Product of the reaction (L-homocysteine) was detected by Ellman’s reagent. Absorbances at 412 nm show no significant difference between *Resc4*-SUMO and buffer control.

To assess cystathionine β-lyase activity, purified SUMO-tagged *Resc4* protein (25 μM) was incubated with 50 μM pyridoxal 5’-phosphate (PLP) and 100 μM L-cystathionine at 37°C for 24 hours. We probed the formation of the L-homocysteine product by reacting with Ellman’s reagent and monitoring absorbance at 412 nm (15). Results showed the absorbance of the sample containing SUMO-tagged *Resc4* was indistinguishable from a control reaction containing no protein (**Figure 2C**). Thus, *Resc4* cannot catalyze cystathionine β-lysis at a detectable level.

### *Resc4* upregulates *genes* in the methionine biosynthesis operon

Since *Resc4* cannot catalyze the enzymatic reaction performed by the deleted natural enzyme, cystathionine β-lyase, we hypothesized that the *de novo* protein may enable the growth of Δ*metC* cells in minimal medium through a regulatory mechanism, as we observed previously for SynSerB3 (7) and SynGltA (8), where the *de novo* protein upregulated expression of a natural enzyme with moonlighting activity for the essential reaction deleted in the auxotrophic strain. To assess this possibility, we probed the transcriptomic and proteomic effects brought about by overexpressing *Resc4* in Δ*metC* cells.

We used RNA-seq to characterize the effect of *Resc4* on the transcriptome of *E. coli* in minimal media. The results showed that *Resc4* upregulates expression of methionine biosynthesis operon in Δ*metC* cells by 30-fold (**Figure 3A, Supplementary Table 2**). One of the overexpressed genes, *metB*, encodes cystathionine γ-synthase (16), which when overexpressed, is known to compensate for the deletion of MetC (17). The upregulation of *metB* by expression of *Resc4* was confirmed by quantitative PCR (fold-change of 8.0 ± 6.4 relative to Δ*metC* cells expressing natural MetC. **Supplementary Figure 4**).

**Figure 3.**
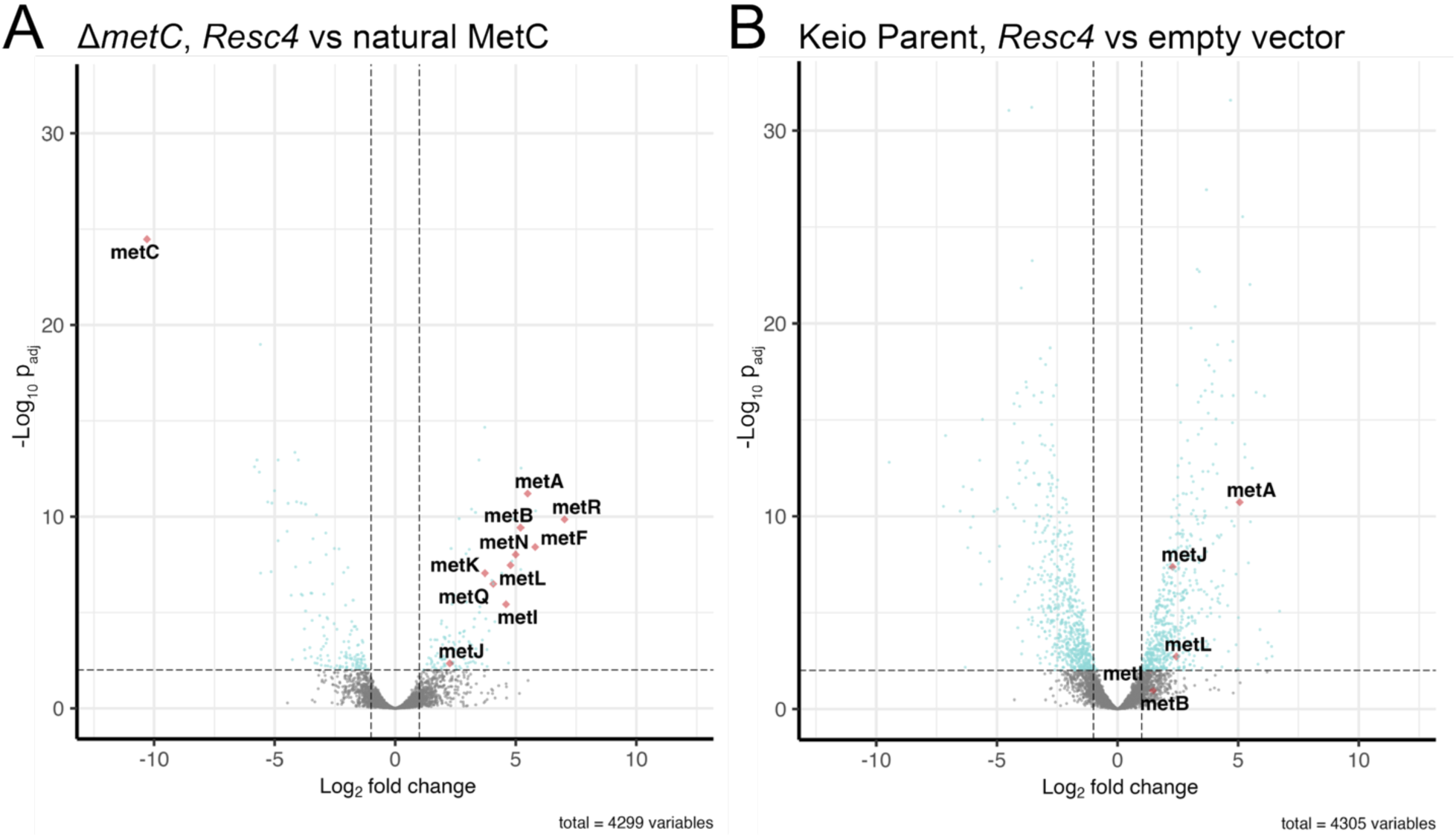
RNA-seq of *Resc4* shows overproduction of met biosynthesis genes and met operon repressor metJ. (A) Transcriptomic effect of *Resc4* relative to natural MetC in Δ*metC* strain. (B) Transcriptomic effect of *Resc4* relative to empty vector in Keio parent strain. Significantly up-regulated or down-regulated met biosynthesis genes are highlighted and labeled.

Since Δ*metC* cells cannot synthesize methionine, we considered the possibility that methionine starvation could in itself lead to increased expression of the Met biosynthesis genes. To control for this possibility, we also probed whether *Resc4* upregulates the methionine biosynthesis operon in a pseudo-wildtype Keio parent strain, where the natural *metC* gene is present in the genome. As shown in **Figure 3B**, even in this pseudo-wildtype background, *metJ*, *metL* and *metA* genes are upregulated by *Resc4* (**Supplementary Table 3**), suggesting the *de novo* protein – not methionine starvation – is responsible for upregulating the Met operon.

### *Resc4* upregulates expression of *proteins* in the methionine biosynthesis pathway

The RNA-seq and qPCR results summarized in the previous section demonstrate that *Resc4* upregulates genes in the methionine biosynthesis operon. To explicitly assess whether the corresponding proteins in this pathway are also overexpressed, we measured the impact of *Resc4* on the proteome of Δ*metC* cells grown in minimal media. These assays showed that proteins in the methionine biosynthesis pathway, including the moonlighting enzyme MetB, are indeed overexpressed in both the soluble and insoluble fraction of the *E. coli* proteome (**Figure 4**).

**Figure 4.**
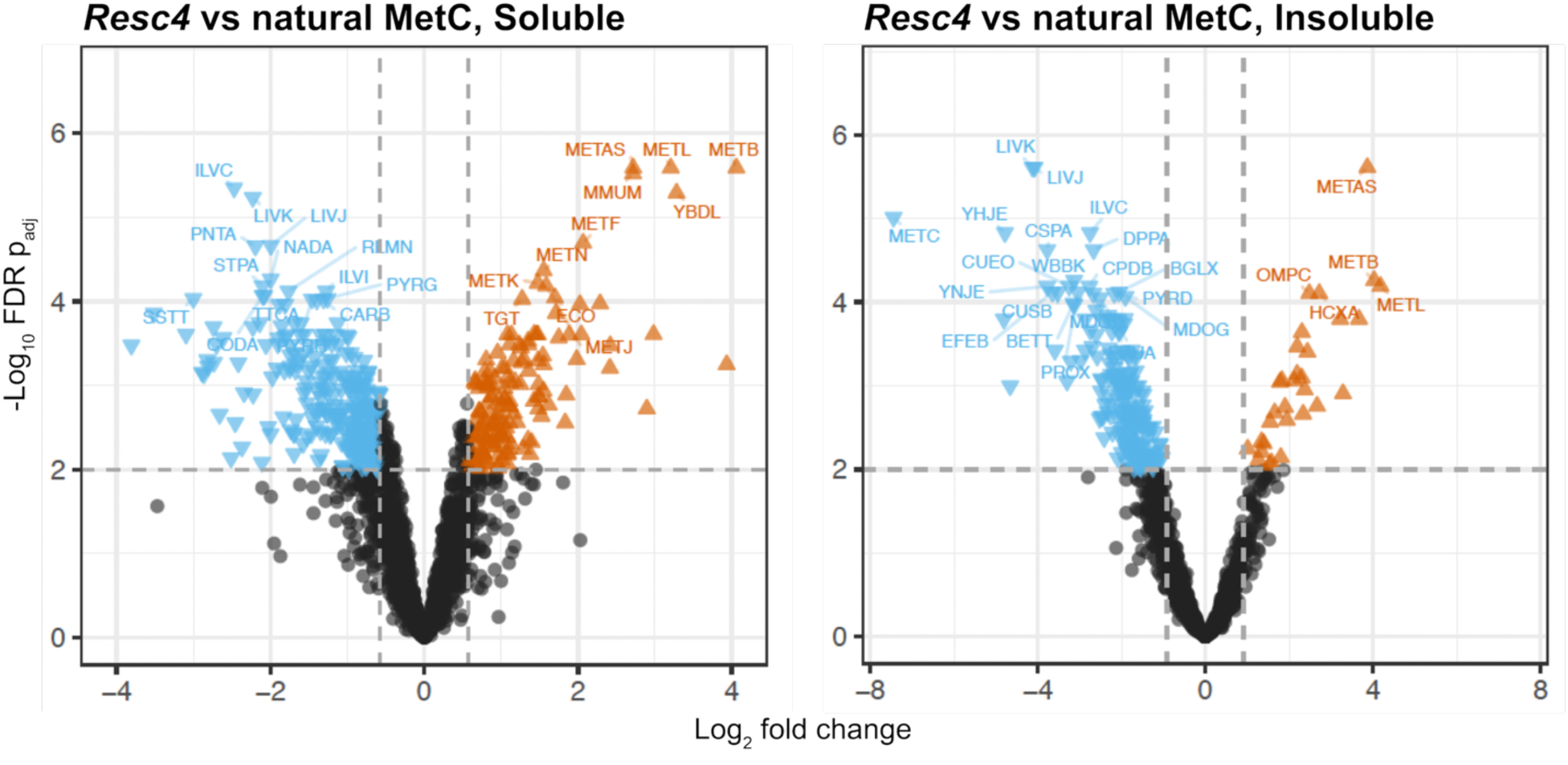
Proteomics analysis of the soluble (left) and insoluble (right) fractions of Δ*metC* expressing *Resc4* relative to natural MetC. Significantly over-expressed proteins are colored in orange, and significantly down-expressed proteins are colored in cyan. Volcano plots are generated by MS-DAP.

The results of these transcriptomic and proteomic studies suggest that *Resc4* rescues the Δ*metC* auxotrophic strain on minimal medium by inducing overexpression of cystathionine γ-synthase, which is encoded by MetB, and known to have a moonlighting activity that compensates for the deletion of cystathionine β-lyase in Δ*metC* cells.

### *Resc4* co-precipitates and sequesters the transcriptional repressor encoded by MetJ

In addition to upregulating the enzymes in the methionine biosynthesis pathway (most importantly cystathionine γ-synthase, encoded by MetB), transcriptomic and proteomic studies revealed that *Resc4* also causes substantial overexpression of the repressor for the Met operon, encoded by MetJ (**Figure 3, 4**) (18). This was surprising because one would typically expect overexpression of a repressor protein to diminish, rather than increase, expression of the corresponding operon. To understand this apparent contradictory behavior, we assessed how *Resc4* impacts the intracellular location (soluble vs. insoluble) of the MetJ encoded repressor protein.

Toward this goal, we measured the interactome of the *Resc4* protein with endogenous *E. coli* proteins. However, because *Resc4* is insoluble, we first had to develop an affinity pulldown assay for an insoluble bait protein. In brief, *Resc4* was dissolved in urea and immobilized on a Ni-NTA column in the presence of urea. The column was then washed with native buffer, and a lysate of soluble proteins from *E. coli* (BL21 DE3 harboring an empty vector) was passed over the column. As a control, the same lysate was also passed over a column of Ni-NTA resin *without* immobilized *Resc4*. After washing the column to remove non-specific binding, the proteins interacting with *Resc4* were eluted with urea and analyzed by timsTOF LC-MS (**Figure 5A**).

**Figure 5.**
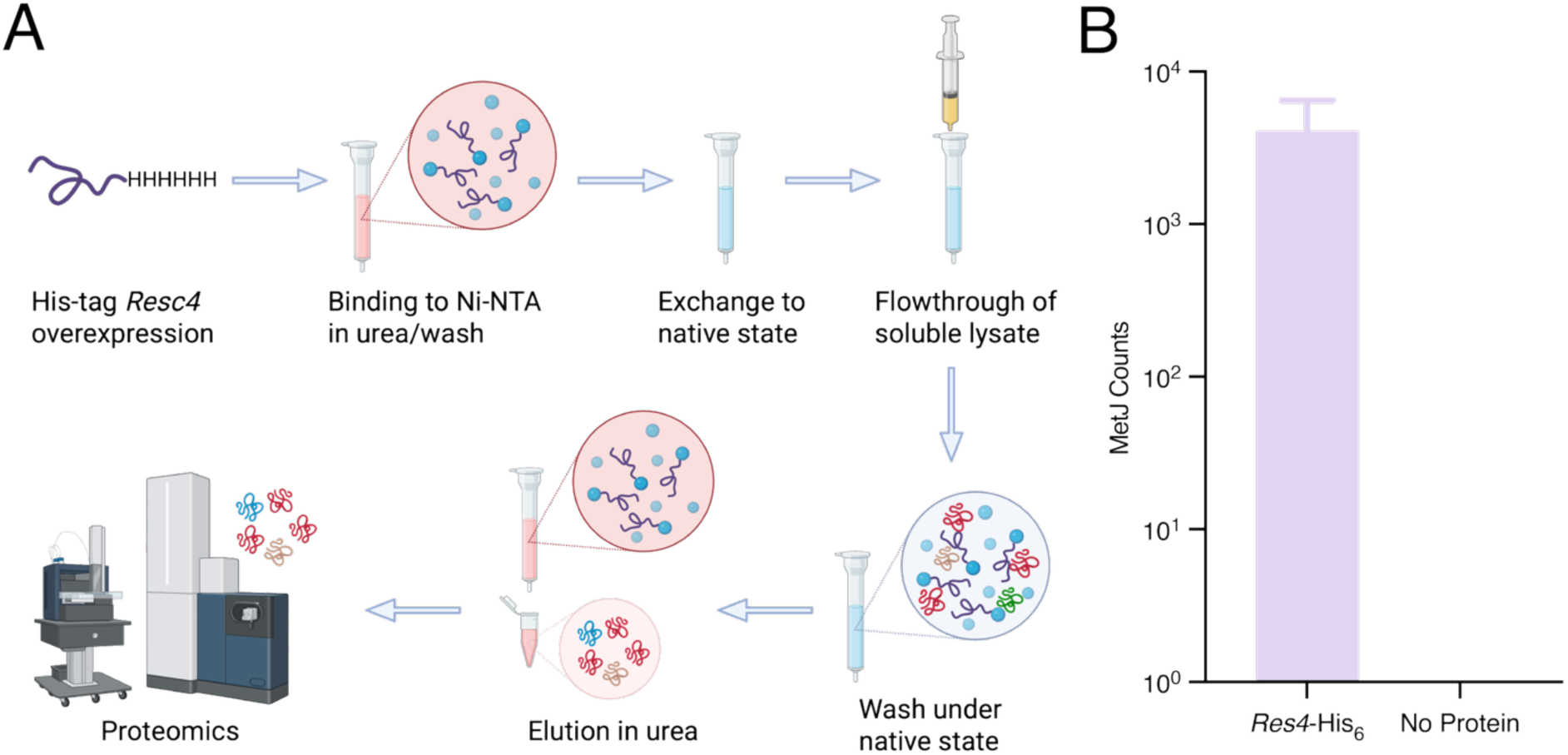
Co-precipitation assay of *Resc4* with *E. coli* soluble proteins. (A) Co-precipitation assay workflow. Created in https://BioRender.com (B) timsTOF-MS analysis of the interactome of *Resc4* with *E. coli* soluble proteins. *Resc4*-His_6_ represents C-terminal His-tagged *Resc4* bound to Ni column. No protein represents no rescuer protein bound to Ni column. No detectable MetJ protein was eluted in the no protein experiment.

Using this technique, we discovered that *Resc4* could bind the MetJ repressor protein, and in the control with no *Resc4* on the column, no bound MetJ repressor was detected (**Figure 5B**). These results suggest that *Resc4* co-precipitates the MetJ repressor protein, sequestering it into the insoluble fraction, and thereby causing upregulation of the methionine biosynthesis operon. Also, *metJ* gene is located in the met operon, where its expression could be repressed by MetJ itself (MetJ is autoregulated) (18). Therefore, sequestration of the MetJ by *Resc4* leads to no active MetJ that can suppress its own expression, which explains *metJ* overexpression under the effect of *Resc4*. After observing experimentally that *Resc4* binds MetJ, we probed this interaction computationally using AlphaFold3 (19, 20). The predicted structure of the *Resc4* monomer suggests the N-terminal a-helix ends after the frameshift, and the C-terminal region of the protein is mostly disordered (**Figure 6A**). Next, we asked AlphaFold3 to predict a structure for the binding of *Resc4* to the MetJ repressor. The four models with the highest prediction confidence suggest that *Resc4* uses its helical region to bind the paired β-strands in the MetJ repressor protein (**Figure 6C**). These are the same paired β-strands that the MetJ repressor uses to bind DNA (**Figure 6B**) (21). These findings suggest that binding *Resc4* blocks the repressor from interacting with DNA, which prevents MetJ from repressing the methionine biosynthetic operon.

**Figure 6:**
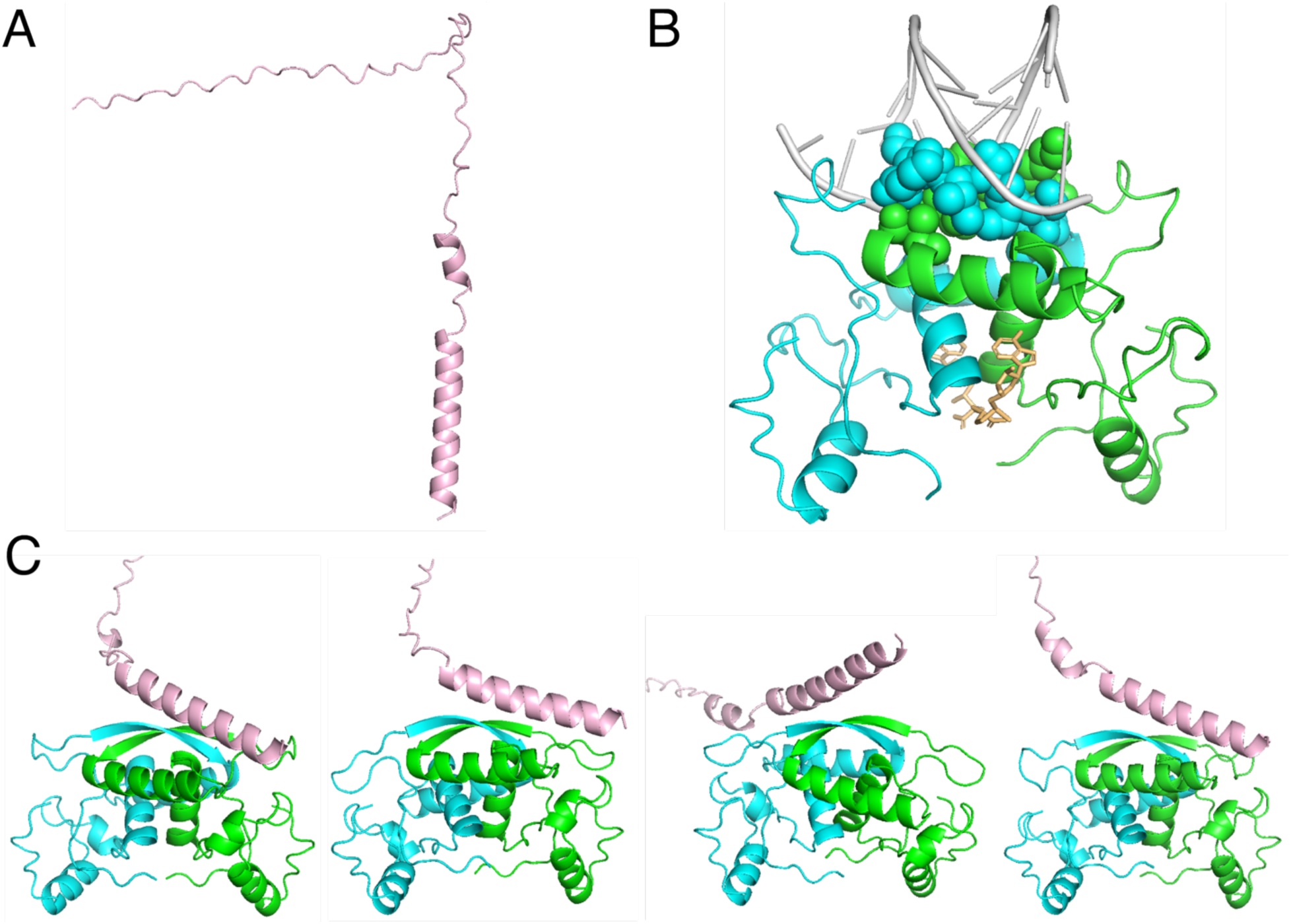
Predicted structure of *Resc4* and MetJ complex. (A) AlphaFold3 prediction of *Resc4*. (B) Crystal structure of MetJ dimer complexed with DNA (PDB ID: 1CMA). Grey: DNA. Brown: S-adenosyl-methionine. Green and cyan: MetJ dimer. Paired β-strands in MetJ that interact with DNA are shown as spheres. (C) AlphaFold3 predictions of *Resc4* and MetJ dimer complex, with 4 highest confidence predictions shown. Green and cyan: MetJ dimer. Pink: *Resc4*. C-terminal loop of *Resc4* is not shown.

### Mutations in key residues of *Resc4* abrogate binding to the repressor encoded by MetJ

The AlphaFold3 predictions highlighted several side chains on *Resc4* that contact the MetJ dimer. In the high-confidence models, Gln7, Glu14, Glu17, Asp18, Gln21, and Ser22 in *Resc4* form polar interactions with the paired β-strands of the MetJ repressor (**Supplementary Figure 5, Supplementary Table 4**). Therefore, we mutated several of these side chains to alternative residues that retained high a-helical propensities (22), but would likely disrupt the intermolecular interactions. We replaced charged residues with the opposite charge to disrupt salt bridge interactions, and uncharged polar residues with Ala to disrupt hydrogen bonding.

We constructed variants E14K, E17K, D18K, Q21A and found that the E14K, E17K and D18K mutations substantially diminished the rescue activity of *Resc4* (**Supplementary Figure 6**). In contrast, the Q21A mutation increased rescue efficiency for reasons that remain unclear.

Overall, the mutagenesis study supports a model wherein *Resc4* co-precipitates and sequesters MetJ repressor protein, thereby preventing it from binding DNA. This leads to overexpression of the methionine biosynthesis enzymes. One of the upregulated enzymes, cystathionine γ-synthase, has a moonlighting activity capable of performing the reaction catalyzed by the cystathionine β-lyase enzyme that was deleted in Δ*metC,* thereby enabling survival of the auxotroph on minimal media.

## Discussion

By screening a library of *de novo* amino acid sequences that share no ancestry with naturally evolved sequences, we discovered a family of frame-shifted and insoluble proteins that enable the growth of the *E. coli* auxotroph Δ*metC* on methionine-deficient media. We purified one of these rescuer proteins, called *Resc4,* and found it has no detectible cystathionine β-lyase activity, thereby suggesting that instead of directly substituting for the deleted natural enzyme, *Resc4* rescues Δ*metC* by an indirect or regulatory mechanism.

Characterization of the transcriptomic, proteomic, and interactomic effects of the *de novo* protein indicates that it enables growth of Δ*metC* by co-precipitating with the natural repressor of the methionine biosynthesis operon, encoded by MetJ. This leads to a 30-fold upregulation of the *metB* gene, which encodes cystathionine γ-synthase (16). This enzyme, when over expressed, can compensate for the deleted activity of cystathionine β-lyase (17), thereby allowing Δ*metC* cells to synthesize methionine and survive on minimal media.

This study adds *Resc4* to the growing list of *de novo* sequences that did not arise in nature, but nonetheless provide life-enabling functions by altering gene regulation. It also highlights that insoluble proteins, which are often considered non-functional or toxic to cells, can enable survival under stressed conditions.

Among the *de novo* life-sustaining proteins that have been reported to date, the binary patterned 4-helix bundle protein, Syn-F4, was shown to rescue the deletion of ferric-enterobactin esterase in Δ*fes* cells through *bona fide* catalytic activity similar to the deleted natural enzyme (5, 6). However, all the other *de novo* life-sustaining sequences characterized to date use a regulatory function to sustain cell growth. For example, several non-natural sequences have been discovered to rescue Δ*serB* (phosphoserine phosphatase) by upregulating HisB, which promiscuously catalyzes phosphoserine dephosphorylation (7, 23). In another example, a different *de novo* 4-helix bundle protein, SynGltA, rescues Δ*gltA* (deleted for citrate synthase) by upregulation of an endogenous enzyme with moonlighting activity that compensates the missing reaction (8). In addition, the Laub lab has shown that screening vast libraries of random sequences led to novel proteins capable of rescuing *E. coli* from the overproduction of MazF toxin by modulating chaperone activity to inactivate the toxin (9). More recently, they showed that a library of random sequences contained novel proteins that use regulatory mechanisms to render *E. coli* resistant to phage infection (10).

Taken together, these findings suggest that in the process of *de novo* gene birth – either in natural evolution or in synthetic biology – new sequences that increase the fitness of an organism may arise more frequently by using regulatory mechanisms rather than direct enzymatic catalysis. As more libraries of novel sequences are screened for additional life sustaining functions, future work – both in our lab and others – will likely reveal myriad ways that novel sequences can enlarge natural proteomes to enhance the fitness of naturally evolved organisms.

## Materials and Methods

### Library construction

We aimed to introduce three distinct degenerate codons at selected positions of the target protein (**Supplementary Figure 1**):

i. **Codon VRM encodes amino acids D, E, H, K, R, N, Q, S, T:** enriched in side chains commonly found in natural enzyme active sites, providing the potential for acid–base catalysis or nucleophilic/electrophilic reactions, thereby generating chemically diverse active centers.
ii. **Codon DTK encodes amino acids F, I, L, V, M:** includes hydrophobic residues of different sizes and shapes, which can modulate cavity volume and geometry and thereby influence substrate binding. The variation in side-chain bulk allows adjustment of the binding pocket’s shape and affinity.
iii. **Codon DNC encodes amino acids N, T, S, I, D, A, G, V, Y, C, F:** covers both polar residues and small hydrophobic residues, which can introduce flexibility and functional diversity while maintaining proper folding stability and avoiding excessive restriction.

We first assembled a Syn-F4 mutation library by PCA-PCR, using the overlapping oligonucleotides listed in **Supplementary Table 5** to introduce degenerate codons at designated sites. Next, to assemble these variants into a single-chain heterodimer library, we linked two Syn-F4 mutants via a flexible peptide (GGGGSGGGGS). Using the Syn-F4 mutation library as template, fragment 1 was amplified with primers F1-F/F1-R and fragment 2 with primers F2-F/F2-R, while the vector backbone was amplified from p3Glar with primers V-F/F-R. Homologous recombination of these three fragments yielded the Syn-F4-Link mutation library, using primers listed in **Supplementary Table 6**.

### Selections for life-sustaining sequences from the heterodimer library

We searched for *E. coli* single gene deletions that are viable in LB but not in M9-glusoce minimal media, according to EcoCyc (24, 25) and Patrick et al (26). From a total of 120 single-gene deletions that fit this criterion, we selected 50 strains based on reaction type and pathway that the deleted-gene is involved in. The strains selected, and the reaction types or biological pathways involved, are listed in **Supplementary Table 1**. The auxotrophy of the selected strains was confirmed by growth on LB plates and non-growth on minimal media plates. In addition to Δ*metC*, we also discovered library sequences that can rescue Δ*fes* and Δ*ilvA*, which are not described in this work.

The chosen strains with single-gene deletions were streaked out from the Keio collection (27) and prepared as electrocompetent cells (ECC). The ECC were then transformed with 20 ng of plasmid library, washed with 1x M9 salts and spread on selective plates. The transformation efficiency was tested by spreading 1:10^5^ dilution of transformed cells on a non-selective plate, where every colony that grew on the non-selective plate indicated 10^5^ transformants. A usual electroporation transformation efficiency was around 10^6^-10^7^ transformants. In addition, to control for cell contamination, the auxotrophic strains transformed with empty vector plasmid or plasmid harboring S824 (28) were spread on a minimal plate with corresponding antibiotics. Competent cells without contamination would lead to a plate with no colonies. As a positive control, the auxotrophic strains transformed with empty vector plasmid or S824 plasmid were spread on minimal media plates supplied with the nutrient the cells cannot synthesize, such as methionine for Δ*metC* cells. The positive control plates all gave a lawn of colonies. All rescue experiments were conducted at 33 °C for 10 days or until colonies were observed on the plates. A workflow of the life-or-death selection process is shown in **Supplementary Figure 2**. The plasmids from the rescued cells were extracted (QIAprep Spin Miniprep Kit). The sequences of the rescuer plasmids were determined by Sanger sequencing (Azenta Life Sciences).

### Enzyme assay for cystathionine β-lysis

To probe the cystathionine β-lyase activity of the rescuer proteins, we overexpressed a rescuer from the screening, *Res4*, in BL21. However, *Resc4* remained in the insoluble fraction of lysed cells. To enhance solubility, we added an His_6_-SUMO tag on the N-terminal of *Resc4* and overexpressed the protein in BL21.

While the SUMO-tagged protein remained insoluble after overexpression, we used 50 mM Na_2_HPO_4_, 300 mM NaCl, 8 M Urea, pH 8.0 to extract the insoluble protein overnight. The solubilized protein was then separated from the insoluble fraction by centrifugation and was passed through a 0.22 μm filter (MilliporeSigma). The urea-solubilized lysate containing overexpressed His_6_-SUMO-*Resc4* was loaded onto a nickel column (HisTrap HP, Cytiva), washed with purification buffer containing 50 mM imidazole, and eluted with 375 mM imidazole in the presence of 8 M urea. The elution fraction was dialyzed sequentially into 4 M urea, 2 M urea and then 0 M urea, and finally filtered (0.22 μm filter, MilliporeSigma). The presence of soluble His_6_-SUMO-*Resc4* was confirmed by SDS-PAGE and mass spectrometry.

25 μM His_6_-SUMO-*Resc4* was incubated with 50 μM PLP and 100 μM L-cystathionine in 50 mM Na_2_HPO_4_, 300 mM NaCl, pH 8.0 buffer at 37 °C. A negative control where no protein was added to the reaction was included. After 24 hours of reaction, Ellman’s reagent was added to the reaction to detect the formation of L-homocysteine using absorbance at 412 nm (15).

### RNA-seq

Δ*metC* or pseudo-wildtype Keio parent *E. coli* cells from the Keio collection (27) harboring either *Resc4*, natural *metC* gene from the ASKA collection (29), or empty vector were grown in M9-glucose minimal media with chloramphenicol and IPTG at 37 °C with shaking. When Δ*metC* cells were used, kanamycin was also included in the culture since Keio collection cells are kanamycin resistant. Cells were collected at OD_600_ 0.2-0.5 and resuspended in 0.5 to 1 mL of DEPC treated water (Invitrogen) and kept frozen as droplets. The total RNA from cells were released from cells by CryoMill and purified by Direct-zol RNA Miniprep kits (Zymo Research). cDNA libraries were prepared by ribo-depletion followed by RNA-Seq Directional Library Prep on Apollo 324 Robot. Next-generation sequencing of the library was done with NovaSeq SP 100nt Lane v1.5 (Princeton University). The sequencing data was processed on Galaxy (30). Specifically, the quality of RNA-seq data was tested by FastQC (31). The data was then processed by Trimmomatic (32) to trim the adapters, mapped to the *E. coli* K12 strain BW25113 reference genome (33) using BWA-MEM (34–36), and the genes counts were determined by htseq-count (37). Differential expression analysis was conducted by DESeq2 in R (38). The volcano plots were created by EnhancedVolcano package in R (39). 3 biological replicates were included for each sample.

### Quantitative PCR (qPCR)

Total RNA from cells was released and purified using the same method as described in RNA-seq. Purified total RNA was then treated with TURBO DNA-free™ Kit (Invitrogen) to remove chromosomal and plasmid DNA. 1 pg of DNA-free total RNA sample was used for cDNA library synthesis using iScript™ cDNA Synthesis Kit (Bio-Rad).

To construct the standard curve for qPCR, 2 pg, 0.2 pg, 0.02 pg, 0.002 pg and 0.0002 pg *metB* amplicon was mixed with qPCR primers (designed by IDT PrimerQuest) and iScript™ cDNA Synthesis Kit (Bio-Rad). The PCR cycle consists of 1 minute of 95 °C denaturation, followed by 42 cycles of amplification (5 seconds at 95 °C, then 30 seconds of annealing/extension at 60 °C). 3 replicates were included for each concentration. The Ct values were used for fitting of the standard curve (**Supplementary Figure 4**).

To quantify the change of *metB* expression in Δ*metC* cells overexpressing *Resc4* relative to natural MetC, 2 biological replicates for each sample, each with 3 technical replicates, were included in the qPCR analysis. After obtaining the Ct value for each sample, the expression levels and fold-change value of *metB* gene were calculated by fitting to the standard curve. The primer sequences used for qPCR are listed in **Supplementary Table 7**.

### Proteomics analysis

Single colonies of Δ*metC* cells, transformed with *Resc4* or natural *metC* plasmids, were used to inoculate LB overnight cultures containing kanamycin, chloramphenicol, and IPTG. The cells from overnight cultures were washed twice with 1 x M9 salts and transferred to 150 mL minimal media with kanamycin, chloramphenicol, and IPTG for growth at 37 °C. Cells from minimal media culture were collected until growth to stationary phase, washed with phosphate-buffered saline (PBS), and stored at −80 °C until cell lysis. To lyse the cells, the cell pellets were resuspended in 10 mL PBS and sonicated for 1 minute 30 seconds. The soluble and insoluble fractions were separated by centrifugation at 35000 x g for 30 minutes. The insoluble proteins were then extracted by 10 mL of 50 mM Na_2_HPO_4_, 300 mM NaCl, 8 M urea, pH 8.0 at 4 °C overnight, and the solubilized proteins were separated from the remaining insoluble fraction by centrifugation at 35000 x g for 1 hour.

To process the soluble and insoluble fractions for proteomics analysis, 50 mM NH_4_HCO_3_, 8 M high-grade urea solution was added to soluble fractions containing 100 μg protein (protein concentration determined by Pierce™ Dilution-Free™ Rapid Gold BCA Protein Assay Kit, Thermo Fisher Scientific) to reach a final urea concentration of 6-8 M. 1 M DTT was added to both soluble and insoluble fractions with a 1:100 v/v ratio to reach a final concentration of 10 mM DTT, where the disulfide bonds were reduced by incubating at 50 °C for 30 minutes. The reactions were then cooled to room temperature, and 300 mM iodoacetamide was added with a ratio of 1:10 to reach a final concentration of 30 mM to covalently modify the reduced cysteine residues. The reactions were incubated at room temperature in dark for 30 minutes to 1 hour. The excess iodoacetamide was neutralized by adding 1:200 1 M DTT and incubating at room temperature for 10 minutes. After incubation, 50 mM NH_4_HCO_3_ was added to the reaction mixture to dilute the urea concentration to less than 2 M, and the protein mixtures were digested with 1:50 w/w trypsin (Trypsin-ultra™, Mass Spectrometry Grade, New England Biolabs) at 37 °C overnight with shaking.

To remove salt and urea from the digested protein samples, Pierce C18 tips (Thermo Fisher Scientific) were used. The tips were primed twice with 200 μL of 40% acetonitrile, 60% water, and 0.1% formic acid. The resin was then activated by 200 μL of 100% water and 0.1% formic acid. To bind peptides to the C18 resin, 200 μL of the digested samples were passed through the resin at a time. The resin was washed twice with 200 μL of 100% water, 0.1% formic acid, and the digested peptides were eluted with 50 μL of 40% acetonitrile, 60% water, 0.1% formic acid.

To further prepare the samples for timsTOF mass spectrometry, the solvents in the eluted peptides were removed by SpeedVac. The peptides were resolubilized in 30 μL of 100% water and 0.1% formic acid and centrifuged at 21300 x g for 1 hour to remove precipitates. The supernatants underwent quality control by QTOF (Agilent) to ensure they did not contain plasticizers.

Proteomics samples were analyzed on timsTOF fleX (Bruker) with PepSep MAX C18 10cm x 150μm, 1.5μm column (Bruker) using data-independent acquisition (DIA). The data was then analyzed using DIA-NN software (40, 41) to obtain protein counts. The fold-change and statistics between different conditions were evaluated using the MS-DAP package in R (42).

### Co-precipitation assay of *Resc4*

To study the interactome of *Resc4*, a His_6_ tag was added to the C-terminal of the rescuer and overexpressed in BL21 DE3. The insoluble protein from overexpression was extracted by PBS with 8 M urea and loaded on 1 mL Ni-NTA resin (Thermo Scientific™ HisPur™ Ni-NTA Resin). The resin was washed twice with 10 mL PBS/8 M urea, twice with 10 mL PBS/8 M urea/50 mM imidazole, and twice with 10 mL PBS/8 M urea. The column was then switched to native condition by passing 10 mL PBS buffer.

To prepare a collection of *E. coli* soluble proteome, BL21 DE3 cells transformed with p3glar empty vector was grown in LB at 37 °C. 100 μM IPTG was added to the culture at OD ∼0.5 and the cells were grown at 18 °C for 18 hours. The cells were collected by centrifugation and resuspended in 30 mL PBS. The cells were lysed by sonication, and the soluble *E. coli* proteome was separated by centrifugation at 35000 x g for 30 minutes followed by filtering.

The soluble lysate of BL21 DE3 with empty vector plasmid was loaded to the Ni-NTA resin with and without *Resc4* bound, and the columns were incubated for 30 minutes to allow binding to the rescuer protein and resin. The resin was then washed twice with 10 mL PBS, twice with 10 mL PBS/50 mM imidazole, and finally twice with 10 mL PBS twice. To elute the proteins that interact with *Resc4* and other non-specific binding, 1 mL 50 mM NH_4_HCO_3_/8 M high-grade urea solution was added to the column. The eluted protein was processed and analyzed using the same method used in the proteomics analysis.

### Mutagenesis of *Resc4*

To remove the methionine start codon in *Resc4*, NEBaseChanger was used to generate the mutation primers. The primer sequences used for methionine start codon deletion are listed in **Supplementary Table 8**.

The residues in *Resc4* that potentially interact with MetJ were identified by structure prediction of *Resc4* in complex with MetJ dimer using AlphaFold Server (20). The top 4 structures with the highest confidence of prediction were analyzed using PyMol (Schrödinger, New York, NY) for interaction residues.

Site-directed mutagenesis was performed using NEBaseChanger (New England Biolabs) or QuikChange (Agilent). Successful incorporation of mutations was confirmed by Sanger sequencing (Azenta Life Sciences). The primer sequences used for site-directed mutagenesis are listed in **Supplementary Table 8**.

## Supplementary Information

**Supplementary Figure 1:**
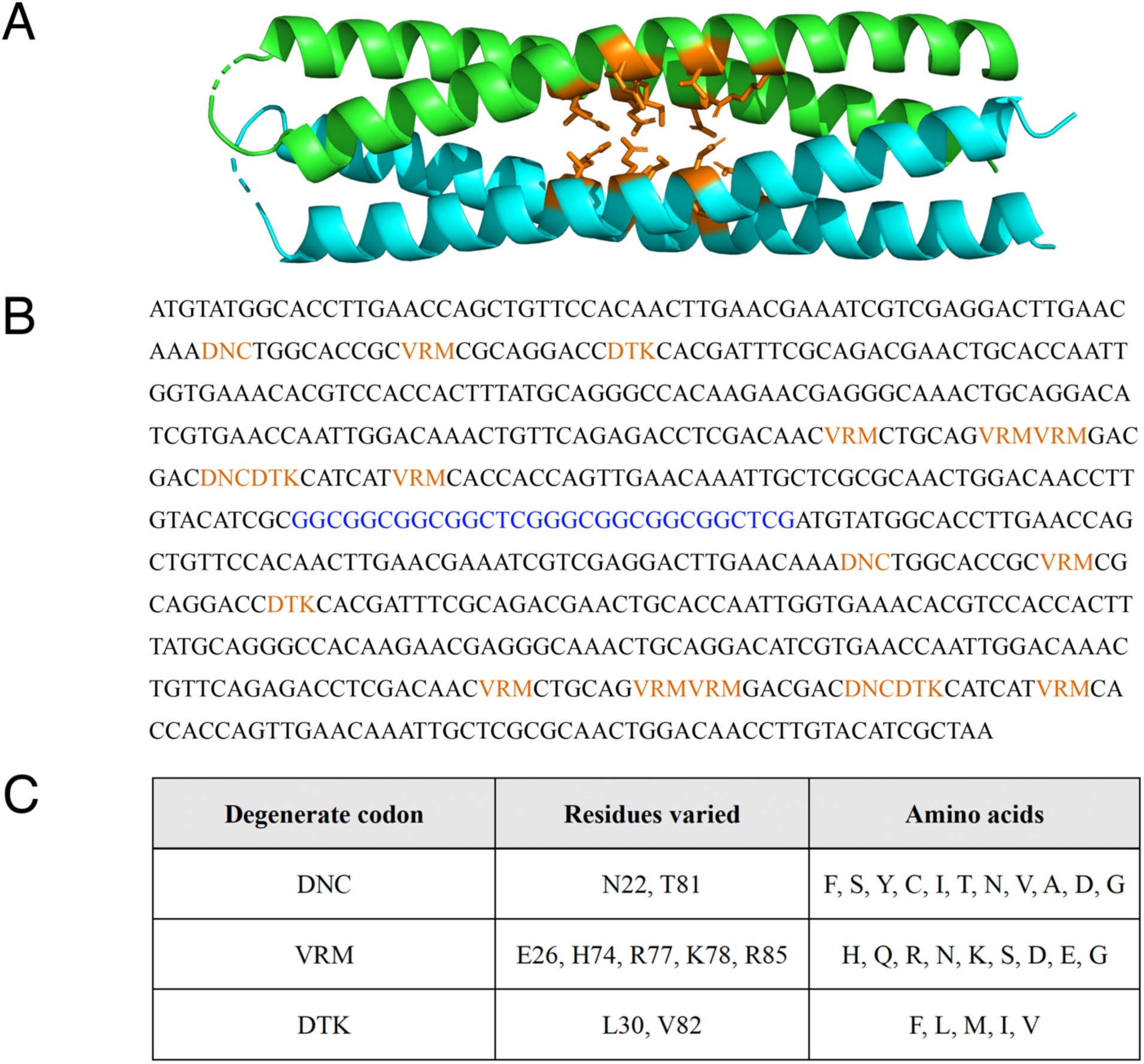
Design of heterodimer library based on Syn-F4-Link. (A) Amino acids that are mutated in the heterodimer library. Side chains of residues that are mutated are labeled in orange. (B) DNA sequence of the heterodimer library. Positions mutated with degenerated codons are labeled in orange. Linker residues are labeled in blue. Abbreviations for degenerate DNA bases are as follows: D: A, G or T. V: A, C or G. R: A or G. M: A or C. K: G or T. N: A, T, C or G. (C) Degenerate codons and corresponding amino acids.

**Supplementary Figure 2:**
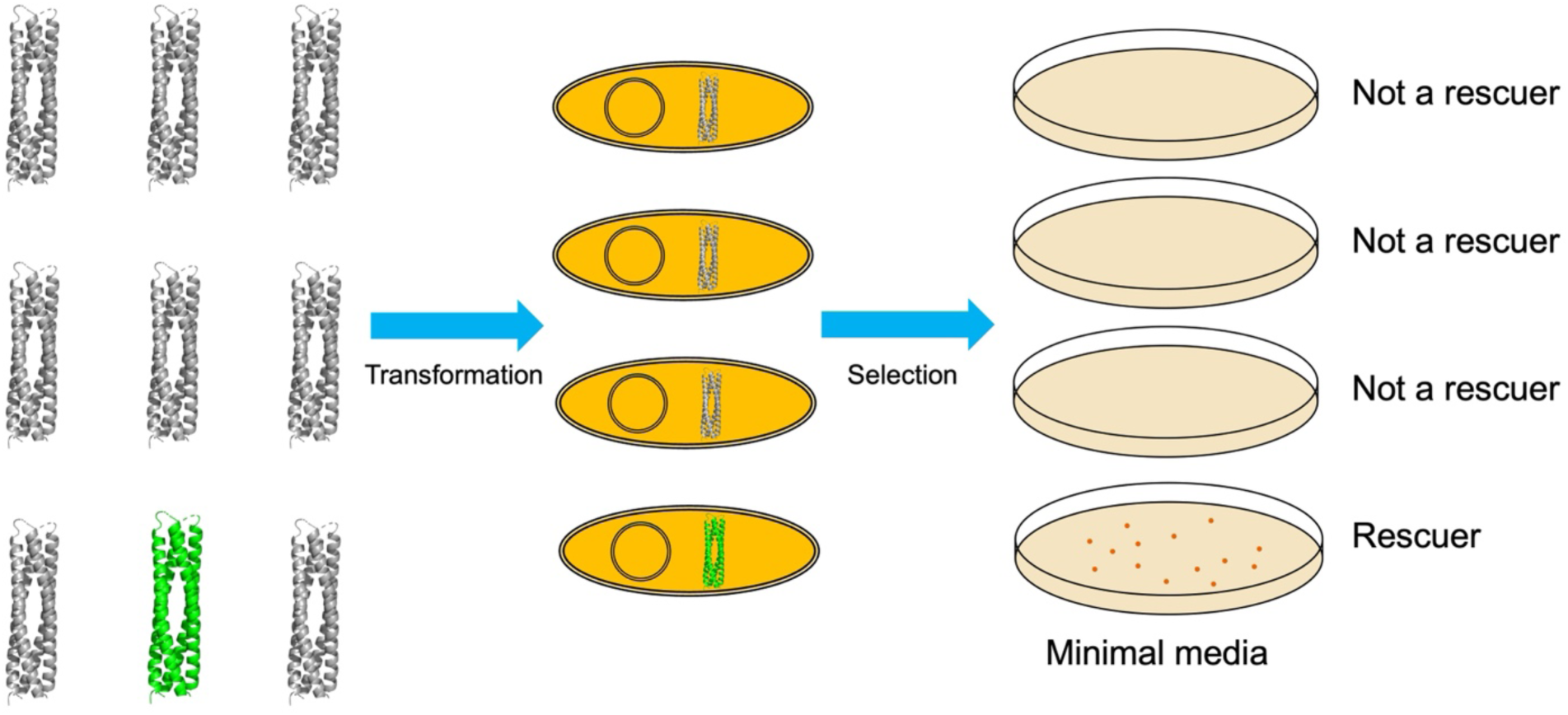
Illustration of auxotroph screening technique.

**Supplementary Figure 3:**
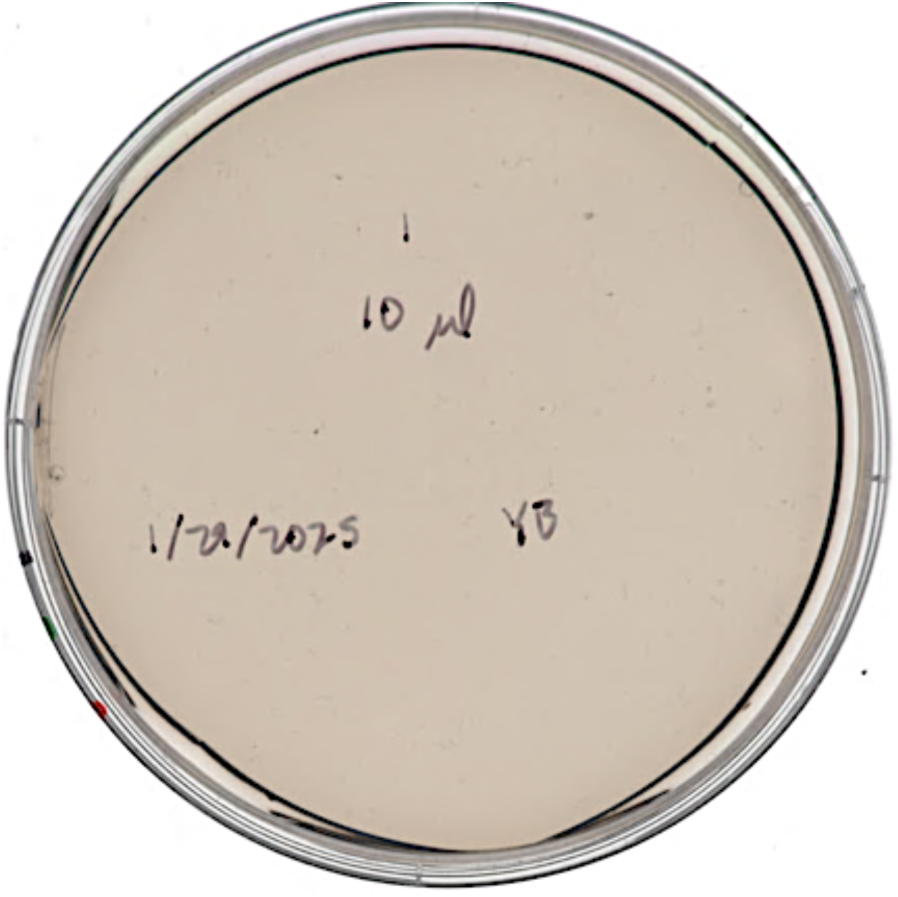
*Resc4* without methionine start codon cannot rescue Δ*metC* on minimal media.

**Supplementary Figure 4:**
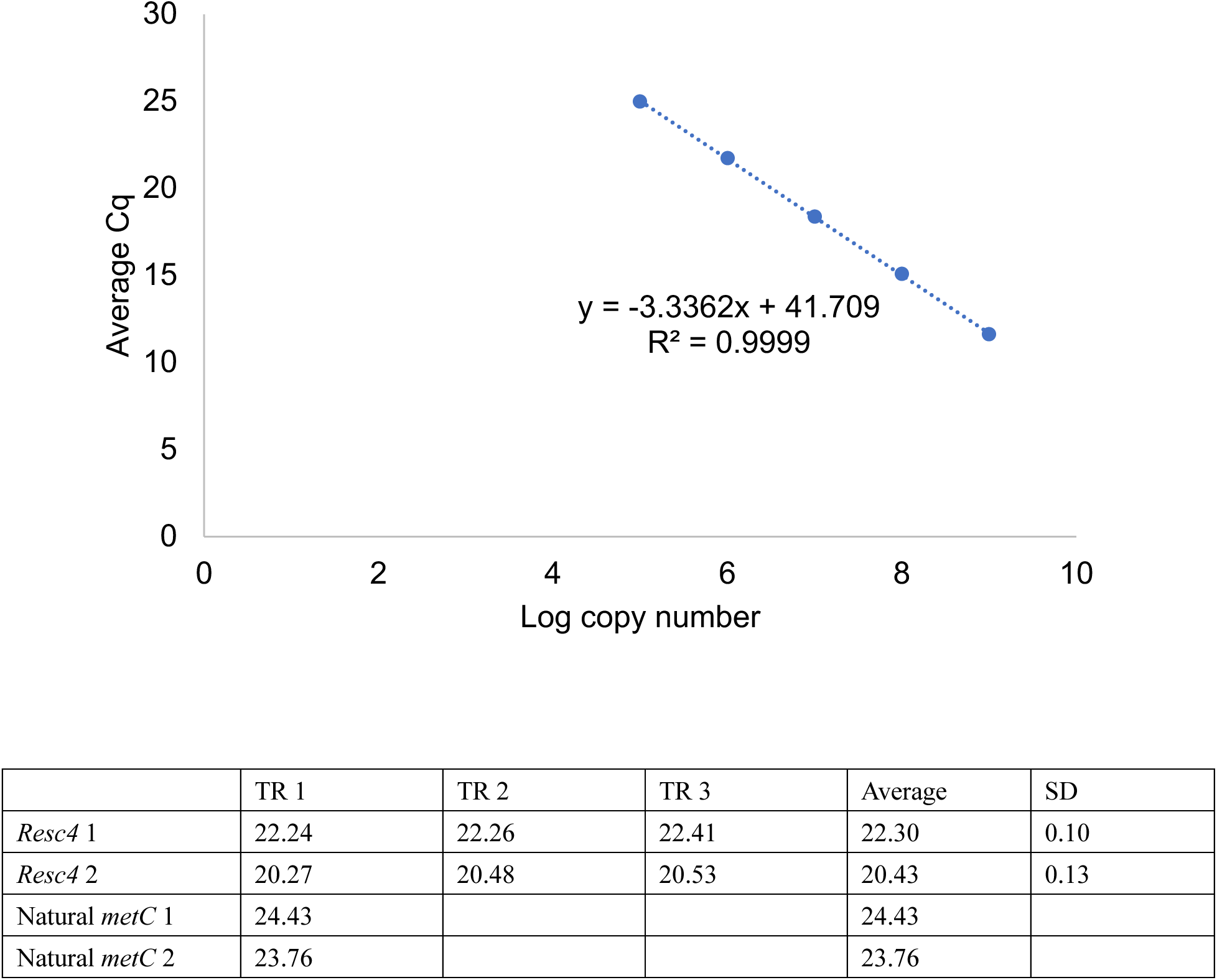
Standard curve and Cq values for qPCR of *metB*. *Resc4* 1/2 and Natural *metC* 1/2 in rows represent biological replicates. TR 1/2/3 in columns represent technical replicates in qPCR.

**Supplementary Figure 5:**
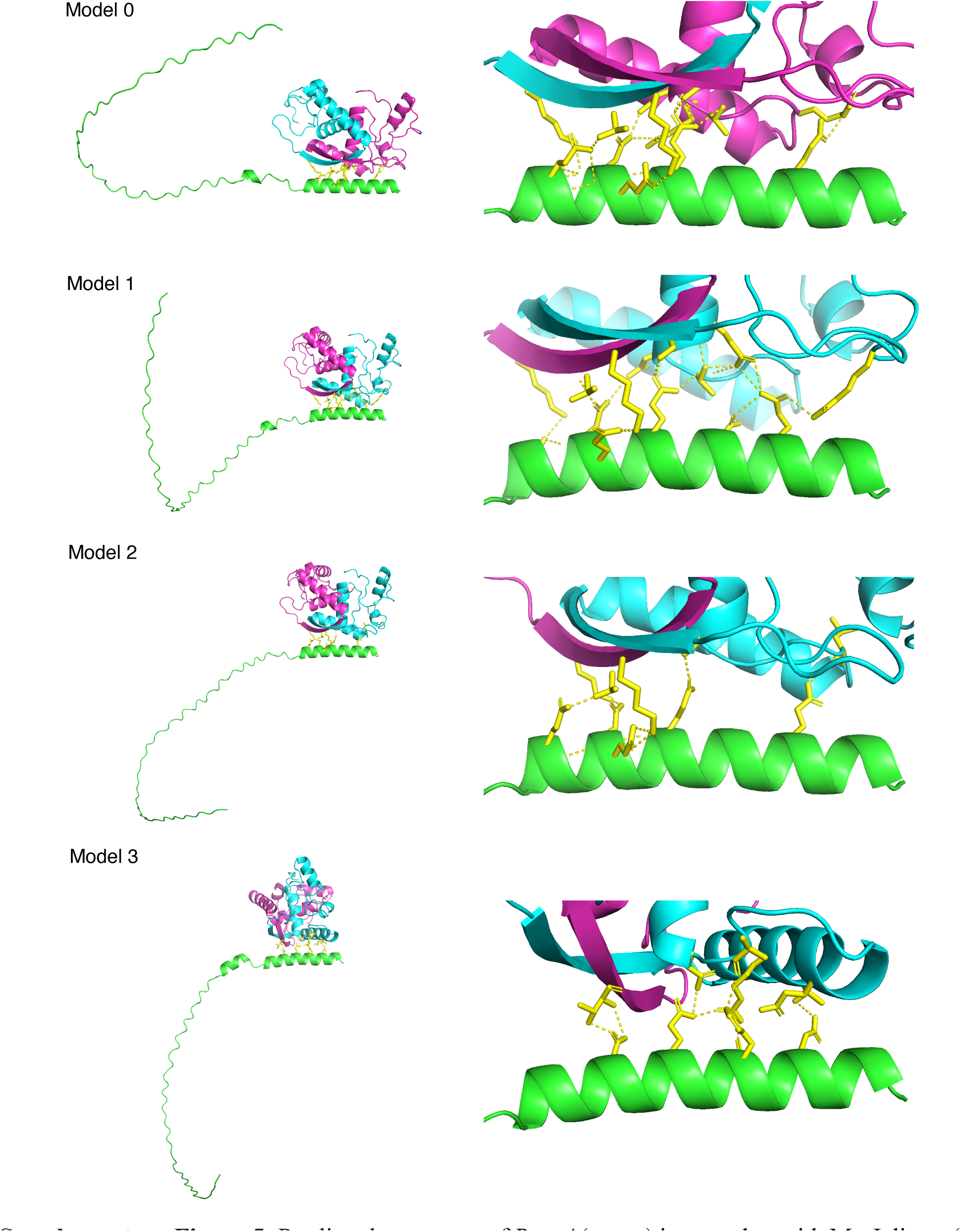
Predicted structures of *Resc4* (green) in complex with MetJ dimer (cyan and magenta). Residues in *Resc4* and MetJ that are in contact are shown in yellow. Polar interactions between residues are indicated in yellow. Model 0 has the highest relative prediction confidence and Model 3 has the lowest relative prediction confidence.

**Supplementary Figure 6:**
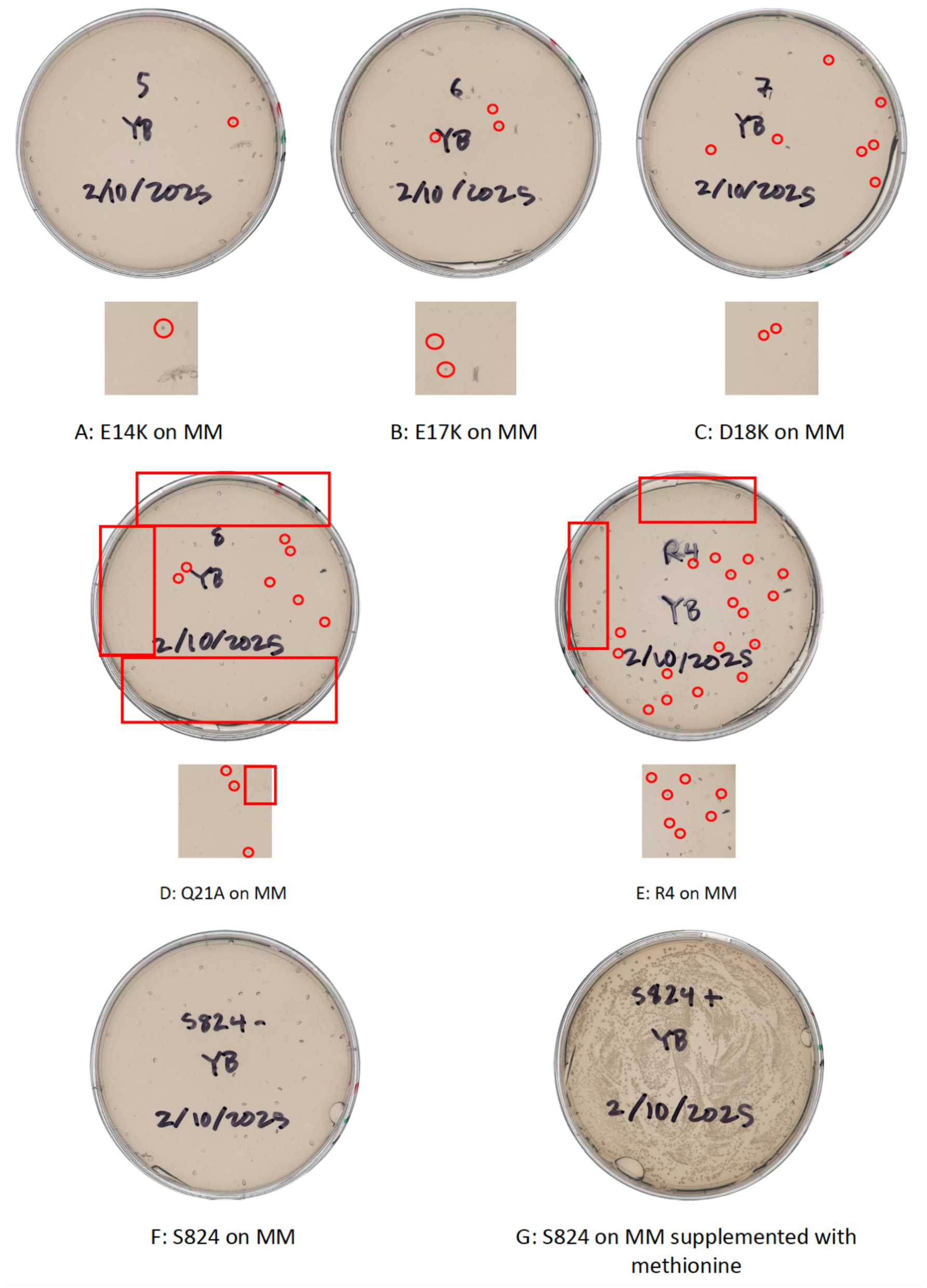
Rescue plates of Δ*metC* by *Resc4* mutants. Red circles and boxes are colonies on agar plate and not bubbles in agar.

**Supplementary Table 1:** Auxotrophic strains used in screening for life-sustaining sequences. The pathways or reaction types that the deleted proteins are involved in are described in the first column.

| Deleted Protein | Strain | Days to Rescue |
| --- | --- | --- |
| Esterase | $\Delta fes$ | 2 days |
| | $\Delta bioH$ | - |
| | $\Delta serB$ | - |
| | $\Delta hisB$ | - |
| | $\Delta cysQ$ | - |
| | $\Delta yjhS$ | - |
| PLP enzyme | $\Delta thrC$ | - |
| | $\Delta hisC$ | - |
| | $\Delta lysA$ | - |
| | $\Delta metB$ | - |
| | $\Delta trpA$ | - |
| | $\Delta trpB$ | - |
| | $\Delta bioA$ | - |
| | $\Delta bioF$ | - |
| | $\Delta iscS$ | - |
| | $\Delta glyA$ | - |
| | $\Delta ilvA$ | 2 days |
| | $\Delta ilvE$ | - |
| | $\Delta metC$ | 5 days |
| | $\Delta serC$ | - |
| Pericyclic | $\Delta pheA$ | - |
| | $\Delta tryA$ | - |
| Arg pathway | $\Delta argA$ | - |
| | $\Delta argB$ | - |
| | $\Delta argC$ | - |
| | $\Delta argE$ | - |
| | $\Delta argG$ | - |
| | $\Delta argH$ | - |
| Aromatic pathway | $\Delta aroA$ | - |
| | $\Delta aroB$ | - |
| | $\Delta aroC$ | - |
| | $\Delta aroD$ | - |
| | $\Delta aroE$ | - |
| Biotin pathway | $\Delta bioB$ | - |
| | $\Delta bioC$ | - |
| | $\Delta bioD$ | - |
| Cys pathway | $\Delta cysC$ | - |
| | $\Delta cysD$ | - |
| | $\Delta cysE$ | - |
| | $\Delta cysH$ | - |
| | $\Delta cysI$ | - |
| | $\Delta cysJ$ | - |
| | $\Delta cysN$ | - |
| Branched amino acid pathway | $\Delta ilvC$ | - |
| Anaerobic respiration | $\Delta gltA$ | - |
| Others | $\Delta cysG$ | - |
| | $\Delta leuD$ | - |
| | $\Delta leuC$ | - |
| Multicopy suppressed | $\Delta glnA$ | - |
| | $\Delta hisH$ | - |
| | $\Delta ilvD$ | - |
| | $\Delta serA$ | - |

**Supplementary Table 2:** Methionine biosynthesis genes in *met* operon with significantly changed expression levels in RNA-seq. The RNA-seq experiment is conducted in Δ*metC* cells.

| **$\Delta metC$** |  |
| --- | --- |
| *Resc4* vs natural MetC |  |
| **Gene** | **log2 foldchange** |
| *metA* | 5.49 |
| *metR* | 7.02 |
| *metB* | 5.20 |
| *metF* | 5.80 |
| *metN* | 5.00 |
| *metL* | 4.78 |
| *metK* | 3.72 |
| *metQ* | 4.07 |
| *metI* | 4.59 |
| *metJ* | 2.26 |

**Supplementary Table 3:** Methionine biosynthesis genes in *met* operon with significantly changed expression levels in RNA-seq. The RNA-seq experiment is conducted in Keio Parent cells where the natural *metC* gene is present.

| Keio Parent |  |
| --- | --- |
| *Resc4* vs empty vector |  |
| Gene | log2 foldchange |
| *metA* | 5.06 |
| *metJ* | 2.29 |
| *metL* | 2.44 |
| *metI* | 1.29 |
| *metH* | 1.22 |
| *metQ* | 1.29 |

**Supplementary Table 4:**
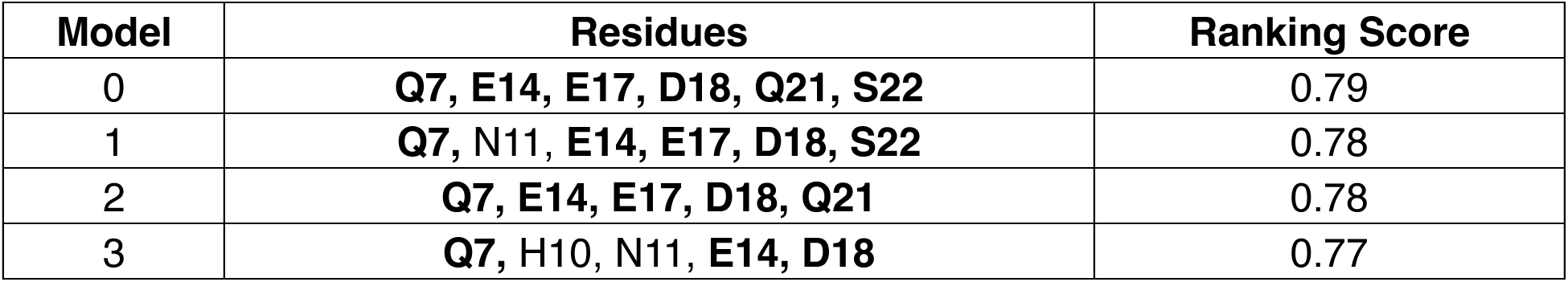
Residues in *Resc4* that interact with MetJ as shown by the top 4 AlphaFold3 prediction models with the highest confidence score. Residues that are selected for mutagenesis are shown in bold.

**Supplementary Table 5:**
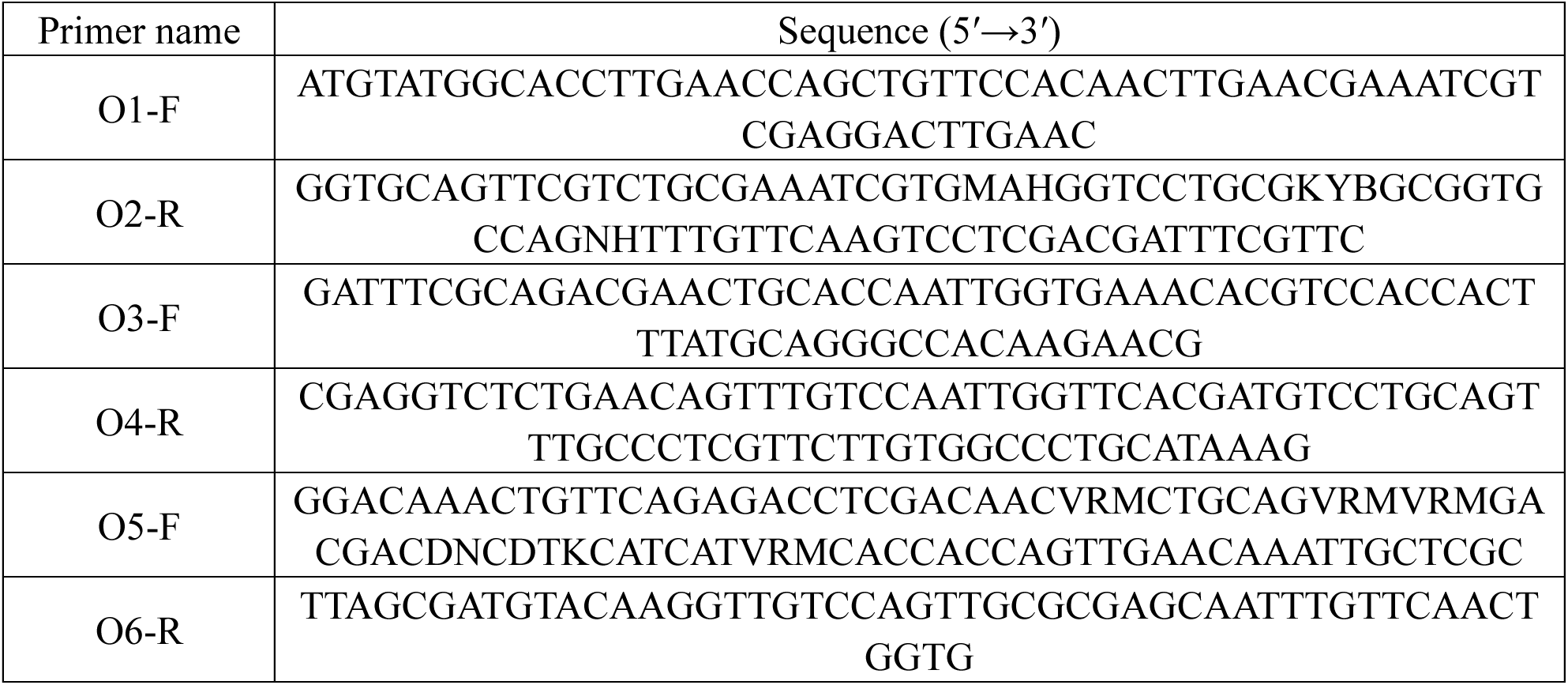
Primers for Syn-F4 mutation library construction.

**Supplementary Table 6:**
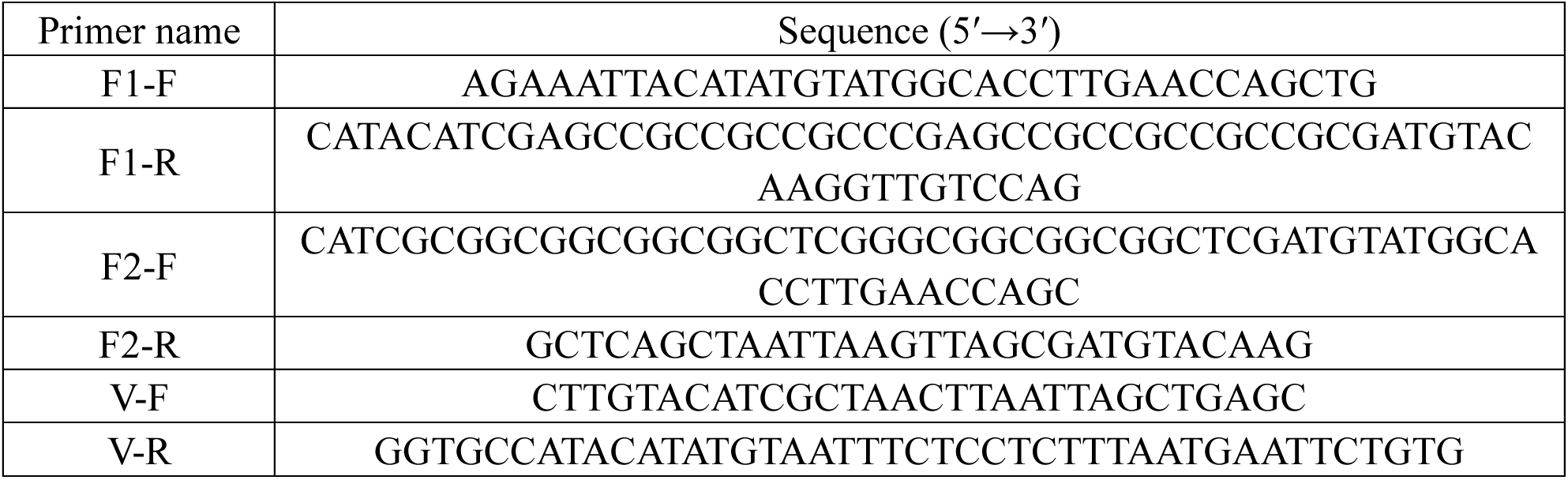
Primers for Syn-F4-Link mutation library construction.

**Supplementary Table 7:**
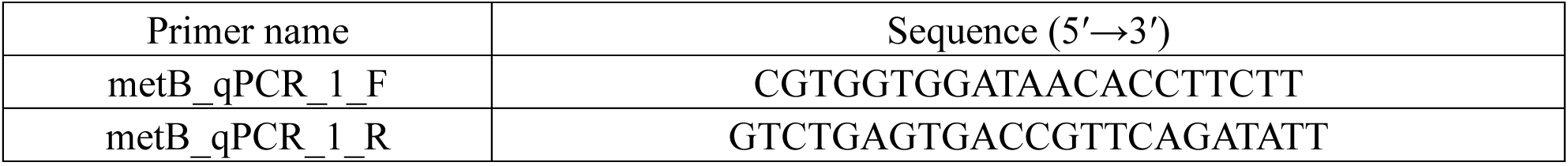
Primers for qPCR analysis of *metB* expression.

**Supplementary Table 8:**
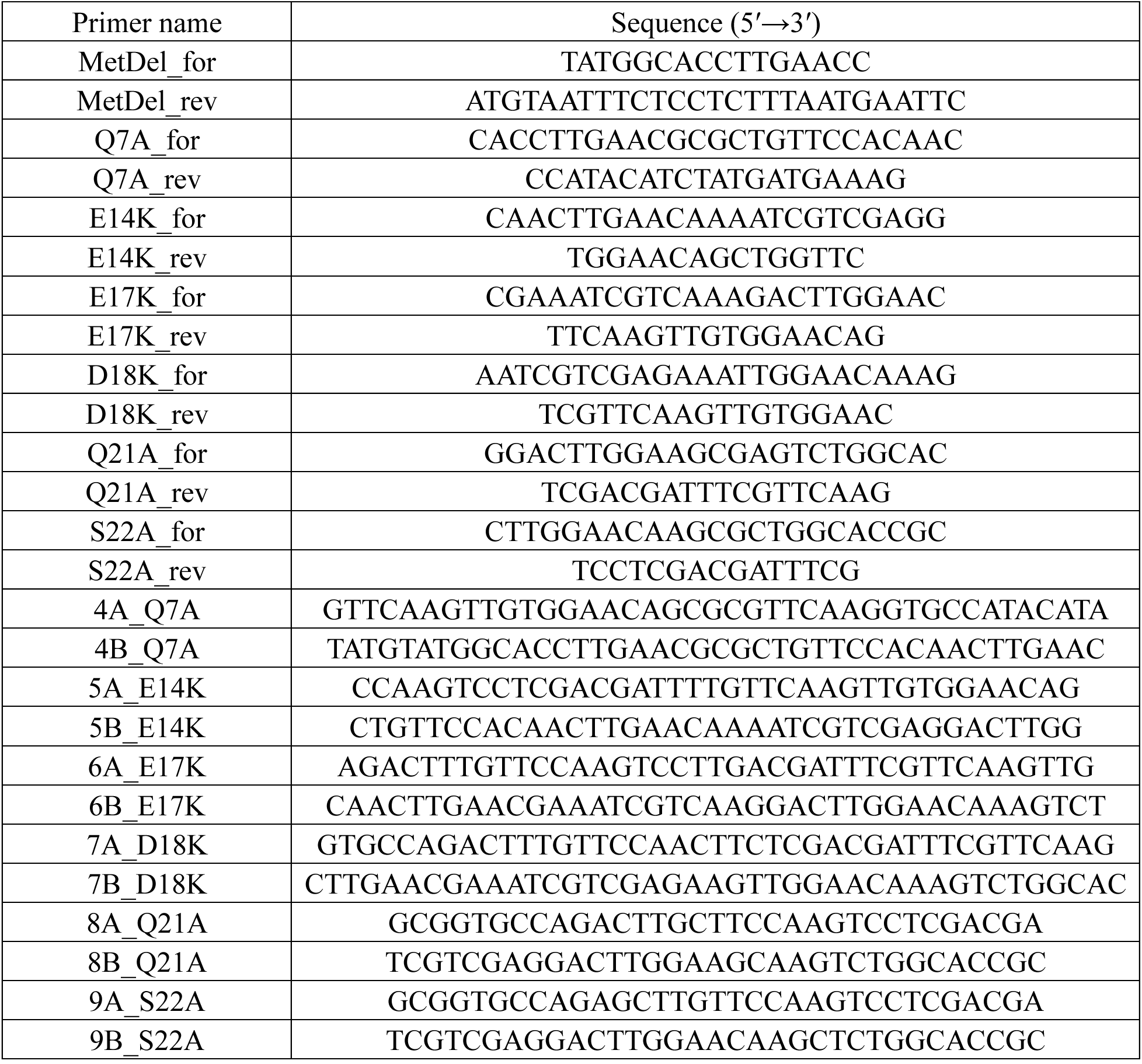
Primers for site-directed mutagenesis of *Resc4*.

## References

1. M. H. Hecht, S. Zarzhitsky, C. Karas, S. Chari, Are natural proteins special? Can we do that? Current Opinion in Structural Biology 48, 124–132 (2018).

2. M. A. Fisher, K. L. McKinley, L. H. Bradley, S. R. Viola, M. H. Hecht, De Novo Designed Proteins from a Library of Artificial Sequences Function in Escherichia Coli and Enable Cell Growth. PLoS ONE 6 (2011).

3. K. J. Hoegler, M. H. Hecht, A de novo protein confers copper resistance in Escherichia coli. Protein Sci 25, 1249–1259 (2016).

4. B. A. Smith, A. E. Mularz, M. H. Hecht, Divergent evolution of a bifunctional de novo protein. Protein Science 24, 246–252 (2015).

5. A. E. Donnelly, G. S. Murphy, K. M. Digianantonio, M. H. Hecht, A de novo enzyme catalyzes a life-sustaining reaction in Escherichia coli. Nature Chemical Biology 14, 253–255 (2018).

6. K. Kurihara et al., Crystal structure and activity of a de novo enzyme, ferric enterobactin esterase Syn-F4. Proc Natl Acad Sci U S A 120, e2218281120 (2023).

7. K. M. Digianantonio, M. H. Hecht, A protein constructed de novo enables cell growth by altering gene regulation. Proceedings of the National Academy of Sciences 113, 2400 (2016).

8. K. M. Digianantonio, M. Korolev, M. H. Hecht, A Non-natural Protein Rescues Cells Deleted for a Key Enzyme in Central Metabolism. ACS Synthetic Biology 6, 694–700 (2017).

9. I. Frumkin, M. T. Laub, Selection of a de novo gene that can promote survival of Escherichia coli by modulating protein homeostasis pathways. Nat Ecol Evol 10.1038/s41559-023-02224-4 (2023).

10. I. Frumkin, C. N. Vassallo, Y. H. Chen, M. T. Laub, Emergence of antiphage functions from random sequence libraries reveals mechanisms of gene birth. Proceedings of the National Academy of Sciences 122, e2513255122 (2025).

11. R. V. Eck, M. O. Dayhoff, Evolution of the Structure of Ferredoxin Based on Living Relics of Primitive Amino Acid Sequences. Science 152, 363 (1966).

12. M. L. Romero Romero, A. Rabin, D. S. Tawfik, Functional Proteins from Short Peptides: Dayhoff’s Hypothesis Turns 50. Angewandte Chemie International Edition 55, 15966–15971 (2016).

13. B. Laber et al., Cloning, purification, and crystallization of Escherichia coli cystathionine beta-lyase. FEBS Lett 379, 94–96 (1996).

14. J. R. Uren, “Cystathionine β-lyase from Escherichia coli” in Methods in Enzymology. (Academic Press, 1987), vol. 143, pp. 483–486.

15. G. L. Ellman, Tissue sulfhydryl groups. Archives of Biochemistry and Biophysics 82, 70–77 (1959).

16. S. V. Tran, E. Schaeffer, O. Bertrand, R. Mariuzza, P. Ferrara, Appendix. Purification, molecular weight, and NH2-terminal sequence of cystathionine gamma-synthase of Escherichia coli. Journal of Biological Chemistry 258, 14872–14873 (1983).

17. M. A. Oberhardt et al., Systems-Wide Prediction of Enzyme Promiscuity Reveals a New Underground Alternative Route for Pyridoxal 5’-Phosphate Production in E. coli. PLOS Computational Biology 12, e1004705 (2016).

18. I. Saint-Girons, N. Duchange, G. N. Cohen, M. M. Zakin, Structure and autoregulation of the metJ regulatory gene in Escherichia coli. Journal of Biological Chemistry 259, 14282–14285 (1984).

19. J. Jumper et al., Highly accurate protein structure prediction with AlphaFold. Nature 596, 583–589 (2021).

20. J. Abramson et al., Accurate structure prediction of biomolecular interactions with AlphaFold 3. Nature 630, 493–500 (2024).

21. J. B. Rafferty, W. S. Somers, I. Saint-Girons, S. E. V. Phillips, Three-dimensional crystal structures of Escherichia coli met repressor with and without corepressor. Nature 341, 705–710 (1989).

22. C. Nick Pace, J. Martin Scholtz, A Helix Propensity Scale Based on Experimental Studies of Peptides and Proteins. Biophysical Journal 75, 422–427 (1998).

23. A. M. Babina et al., Rescue of Escherichia coli auxotrophy by de novo small proteins. eLife 12, e78299 (2023).

24. D. Karp Peter et al., The EcoCyc Database (2023). EcoSal Plus 11, eesp-0002-2023 (2023).

25. Lisa R. Moore et al., Revisiting the y-ome of Escherichia coli. Nucleic Acids Research 52, 12201–12207 (2024).

26. W. M. Patrick, E. M. Quandt, D. B. Swartzlander, I. Matsumura, Multicopy Suppression Underpins Metabolic Evolvability. Molecular Biology and Evolution 24, 2716–2722 (2007).

27. T. Baba et al., Construction of Escherichia coli K-12 in-frame, single-gene knockout mutants: the Keio collection. Molecular Systems Biology 2, 2006.0008 (2006).

28. Y. Wei, S. Kim, D. Fela, J. Baum, M. H. Hecht, Solution structure of a de novo protein from a designed combinatorial library. Proceedings of the National Academy of Sciences 100, 13270 (2003).

29. M. Kitagawa et al., Complete set of ORF clones of Escherichia coli ASKA library ( A Complete S et of E. coli K −12 ORF Archive): Unique Resources for Biological Research. DNA Research 12, 291–299 (2005).

30. T. G. Community, The Galaxy platform for accessible, reproducible, and collaborative data analyses: 2024 update. Nucleic Acids Research 52, W83–W94 (2024).

31. S. Andrews (2010) FastQC: A quality control tool for high throughput sequence data.

32. A. M. Bolger, M. Lohse, B. Usadel, Trimmomatic: a flexible trimmer for Illumina sequence data. Bioinformatics 30, 2114–2120 (2014).

33. F. Grenier, D. Matteau, V. Baby, S. Rodrigue, Complete Genome Sequence of Escherichia coli BW25113. Genome Announcements 2, 10.1128/genomea.01038-01014 (2014).

34. H. Li, R. Durbin, Fast and accurate short read alignment with Burrows–Wheeler transform. Bioinformatics 25, 1754–1760 (2009).

35. H. Li, R. Durbin, Fast and accurate long-read alignment with Burrows–Wheeler transform. Bioinformatics 26, 589–595 (2010).

36. H. Li, Aligning sequence reads, clone sequences and assembly contigs with BWA-MEM. arXiv preprint arXiv:1303.3997 (2013).

37. S. Anders, P. T. Pyl, W. Huber, HTSeq—a Python framework to work with high-throughput sequencing data. Bioinformatics 31, 166–169 (2014).

38. M. I. Love, W. Huber, S. Anders, Moderated estimation of fold change and dispersion for RNA-seq data with DESeq2. Genome Biology 15, 550 (2014).

39. K. Blighe, S. Rana, E. Turkes (2024) EnhancedVolcano: Publication-ready volcano plots with enhanced colouring and labeling. R package version 1.24.0.

40. V. Demichev, C. B. Messner, S. I. Vernardis, K. S. Lilley, M. Ralser, DIA-NN: neural networks and interference correction enable deep proteome coverage in high throughput. Nature Methods 17, 41–44 (2020).

41. V. Demichev et al., dia-PASEF data analysis using FragPipe and DIA-NN for deep proteomics of low sample amounts. Nature Communications 13, 3944 (2022).

42. F. Koopmans, K. W. Li, R. V. Klaassen, A. B. Smit, MS-DAP Platform for Downstream Data Analysis of Label-Free Proteomics Uncovers Optimal Workflows in Benchmark Data Sets and Increased Sensitivity in Analysis of Alzheimer’s Biomarker Data. Journal of Proteome Research 22, 374–386 (2023).

